# Characterization of wheat lacking B-type starch granules

**DOI:** 10.1101/2021.10.01.462759

**Authors:** Benedetta Saccomanno, Pierre Berbezy, Kim Findlay, Jennifer Shoesmith, Cristobal Uauy, Bruno Viallis, Kay Trafford

**Affiliations:** The National Institute of Agricultural Botany, Huntingdon Road, Cambridge, Cambridgeshire CB3 0LE, UK; Limagrain Céréale Ingrédients, ZAC les Porte de Riom, Avenue George Gershwin, 63200 RIOM, France; The John Innes Centre, Norwich Research Park, Colney Lane, Norwich, Norfolk NR4 7UH, UK

**Keywords:** B-granule-less, BlessT, bread wheat, starch granule, wheat grain, wheat flour

## Abstract

The physicochemical and agronomic properties of a new form of bread wheat, lacking B-type starch granules (BlessT) were assessed. Three BlessT mutant lines, made by combining homoeologous deletions of *BGC1*, a gene responsible for the control of B-granule content were compared with two sibling lines with normal starch phenotype and the parent line, cv. Paragon. Quantification of starch granule size and number in developing grain confirmed the lack of small, B-type starch granules throughout development in BlessT. Most starch, flour, grain and loaf characteristics did not vary between BlessT and the wild type sibling controls. However, BlessT starches had higher water absorption, reduced grain hardness and higher protein content, and dough made from BlessT flour required more water and had increased elasticity. Despite the lack of B-granules, BlessT lines do not display a significant decrease in total starch content suggesting that it should be possible to produce commercial wheat varieties that lack B-type starch granules without compromising yield. These findings support the potential utility of this novel type of wheat as a specialist crop in applications ranging from bread making and alcohol production to improved industrial starch products.

## 1. Introduction

The functionality of cereal grain, flour and starch is influenced by the shape and size of the starch granules within the endosperm. Two types of granules, (A- and B-type), exist in the grains of bread wheat (*Triticum aestivum. L*.) (and other Triticeae species such as *Triticum durum L*., *Hordeum vulgare L*. and *Secale cereale L*.). Compared to the A-type, B-type starch granules are far more numerous, smaller, more spherical in shape and they initiate approximately ten days later than the A-type during grain development (Parker, 1985; Bechtel et al., 1990). B-granules have several negative impacts on end-uses: half of the B-granules are broken down during barley malting (representing a loss of 5% of total starch) (Tillett and Bryce, 1993); barley B-granules cause a ‘starch haze’ that can cause filtration problems during brewing (Bathgate et al., 1974); and B-granules are easily lost in the waste stream during starch purification (Stoddard and Sarker, 2000). However, B-granules have a positive impact on the functional properties of pasta (Soh et al., 2006) and may be required for optimal loaf volume (Soulaka and Morrison 1985; Park et al., 2009).

In the past, the lack of sufficient natural variation in starch granule size distribution amongst domesticated Triticeae species made it impossible to breed cultivars lacking B-granules. However, a few wild relatives of wheat (goat-grasses) naturally lack B-granules, including *Aegilops peregrina* (Stoddard and Sarker, 2000). By generating and analysing a population of Aegilops segregating for B-granule number, we identified a locus, *BGC1* responsible for controlling B-granule content (Howard *et al*., 2011). We then selected mutant lines of bread wheat (cv. Paragon, wild type *BGC1* genotype: AABBDD) with deletions encompassing the orthologous *BGC1* loci (Chia *et al*., 2017). Single mutant plants with homozygous 4A or 4D deletions (--BBDD and AABB--, respectively) were crossed together and progeny plants homozygous for both deletions were selected from the resulting F_2_ and F_3_ progeny (--BB--double mutants). From the same progeny lines, we also selected wild type sibling lines that were homozygous wild type for both the A and D genome *BGC1* mutations (AABBDD). In contrast to the parent line, Paragon and the single deletion plants (which had normal B-granule content), double deletion mutant plants were found to lack B-type starch granules. Subsequent work using wheat TILLING mutants of tetraploid and hexaploid wheat confirmed the identity of the *BGC1* gene (Chia *et al*., 2020). Small scale assays of glasshouse grown plants showed that the B-less wheat grains had normal A-type starch granule morphology, normal overall starch content, and normal grain weight (Chia *et al*., 2017). In addition to variation in starch granule-size distribution, the B-less wheat grains differed from controls in grain hardness, starch swelling power, and amylose content. The new types of wheat (Paragon AD double-deletion mutants) were named BlessT (B-less Triticum).

The *BGC1* 4A and 4D deletions each span ∼600–700 genes and both of the single mutant BlessT parent lines had multiple other large deletions present in their genomes (Chia *et al*., 2017). Each of the double mutant and sibling wild type lines will have inherited an unknown combination of these additional unlinked deletions. To further test the effects of the lack of B-type starch granules on grain, flour and starch functional properties, we chose three independently selected mutant lines (MUT1, MUT4, MUT23) and two wild type sibling lines with normal starch phenotype (WT1 and WT4) and these were grown, together with the wild type parent line (Paragon) in the field and glasshouse over two successive years. Comparison of the wild type sibling lines with wild type Paragon was used to assess the effects of the unlinked (non-*BGC1*) deletions. Comparison of the three BlessT mutant lines was used to determine effects common to all three mutants (and therefore likely to be due to deletion of *BGC1* or the adjacent genes). Starch granule size during grain development as well as plant growth, yield parameters, and the functional properties of starch and flour were assessed.

## 2. Materials and methods

### 2.1. Germplasm

The BlessT deletion mutant lines (MUT1, MUT4, MUT23) and wild-type sibling controls (WT1, WT4) were selected from the F_2_ or F_3_ progeny of crosses between the same two single-deletion mutants (A1 and D4) of cv. Paragon (Chia *et al*., 2017). The pedigree of these lines is shown in Supplementary Figure S2.

### 2.2. Plant growth

Plants were sown in a field in Cambridge (location decimal degrees 52.240704, 0.099924946) in the spring of 2018 in plots that were either large (2 × 4 m) or small (1 × 1 m). All plots were supplied with a comprehensive agronomy regime that included nitrogen application before drilling, herbicides and fungicides. Triplicated, spatially-randomised large plots were used for measurement of growth metrics and yield parameters. Non-randomized large plots were used for bulking of grain samples. The small plots were used for analysis of developing grains and for preparation of starch and flour for physicochemical analyses at NIAB, UK. Subsequently, the remaining grains of each genotype were combined (pooled samples) and those of WT1, WT4, MUT4 and MUT23 were used for additional assessment of functional properties, including baking trials, at Limagrain Céréale Ingrédients (LCI), France. The plants grown in the field in 2018 were found to have high α-amylase activity. This is frequently caused by excess rainfall at or near harvest and it can impair bread-making quality. Some tests (Hagberg falling number, granule-size distribution and RVA) were therefore also carried out at LCI on material that had been grown in the ground in an unlit glasshouse in Cambridge in the summer of 2017. These plants were provided with fertilizer and were watered only as required.

### 2.3. Starch granule size and number in developing grain

The primary tillers were tagged at Zadoks growth stage 59 (ear fully emerged) and developmental stage was measured as days after this stage. Developing grains were harvested at various stages between 6 to 40 days after emergence and each sample included 2 grains per spike from 3 or 6 different plants. Starch was extracted as described in South and Morrison (1990) except that, to improve the purity of the starch, the sodium chloride/sucrose washing solution was replaced with 80% (w/v) CsCl.

Starch granule number per endosperm was measured using an automated image analysis method. Dry starch was suspended in 50-200 μl of water to give an approximate concentration of 10 ×10^6^ granules per millilitre. Duplicate samples were loaded onto slides and both granule-size distribution and number per unit volume was measured using a Biorad TC20 cell counter. The values for small granules with a diameter of 10 μm and below (mainly B-granules) was compared with those of granules with diameters greater than 10 μm (mainly A-granules). Starch granule-size distribution was also measured using a Coulter counter as reported previously (Chia *et al*., 2020).

### 2.4. Scanning electron microscopy (SEM)

Drops of purified starch grains were dried overnight on a glass coverslip which was mounted on the surface of an aluminium pin stub using a double-sided adhesive carbon disc (Agar Scientific Ltd, Stansted, UK). The stubs were then sputter coated with approximately 15 nm of gold in a high-resolution sputter-coater (Agar Scientific Ltd) and transferred to a FEI Nova NanoSEM 450 FEG scanning electron microscope (FEI, Eindhoven, The Netherlands). The samples were viewed at 3 kV and digital TIFF files were stored.

### 2.5. Growth metrics

For all measurements, plants at the edges of the plots were excluded. Establishment rate was assessed two weeks after drilling as number of plants on both sides of a 1-meter ruler. Plant height was measured using a ruler from the ground to the tip of the ear 10 weeks after drilling. Flowering time was scored as days from drilling to Zadoks growth stage 59 (ear fully emerged) rather than days to anthesis since this can be scored more accurately. Tiller count was assessed 12 weeks after drilling as the number of tillers in a 1-m^2^ quadrat. Plant density was assessed as NDVI measurement throughout the growing season using a RapidSCAN CS-45, however, only representative measurements taken midseason are presented here.

### 2.6. Yield parameters

Grain width, length, area and thousand grain weight (ie. average grain weight x 1000) of a random subsample of 600-800 grains per plot (four replicates, each from a different plot) were measured using a MARVIN seed analyser 5.0 (GTA Sensorik, Neubrandenburg, Germany), according to the manufacturer’s instructions. Measurements of plot yield and hectolitre weight were made during harvesting, on combine (Haldrup C-85 plot combine).

### 2.7. Grain characterization

Protein and moisture content were measured (with technical duplication) on four replicate samples of ∼200 g of grain (each from a separate small plot) using a FOSS 6500 wavelength-scanning, near-infrared (NIR) spectroscope (incorporating ISIScan software) according to the manufacturer’s instructions. The sample spectra were compared with calibration spectra taken from wheat grain samples with known protein, and moisture contents. Grain hardness was determined for four replicate samples per genotype (each from a different small plot) using a single kernel characterization system (Perten SKCS4100) according to the manufacturer’s instructions. Each sample consisted of 20 replicate grains.

For measurement of germination rate, 100 grains drawn at random were placed onto two sheets of germination paper and covered by a third sheet. Prior to use, each sheet was soaked in 22 ml of water. Grains were incubated for three days at 10°C and then 4 days at 20°C before assessment. Seed viability was assessed for 100 grains selected at random. Grains were soaked in water for a minimum of two hours at 40°C. The embryo was then removed from the grain and incubated in a 1% (w/v) solution of 2,3,5-triphenyl tetrazolium chloride overnight at 30°C. Embryos were considered inviable if the shoot or the scutellum or all three roots were unstained.

### 2.8. Flour characterization

Flour was prepared from four replicate 50-g samples of grains (each from a separate small plot) by milling in a Foss Cyclotec 1093 mill with 1 mm screen (plot samples) or using a Buhler 202 mill (pooled samples). Hagberg Falling number was measured with a Perten 1700. Flour (7g) was mixed with 25 ml of water in a glass test tube and the time taken by the plunger to reach the bottom of the tube was measured. Alpha-amylase activity was measured using Megazyme K-AMYL SD Kit, according to manufacturer’s instructions (plot samples) or activity was inferred from the RVA data (pooled samples).

### 2.9. Flour Rheology

All measurements were performed on pooled samples using manufacturer’s recommended methods as follows. Alveograph (Chopin): Norm NF EN ISO 27971. Farinograph (Brabender): Norm AFNOR NF ISO 5530-1. Rapid visco analysis (RVA) (Perten): standard protocol 50-95-50 °C (with and without silver nitrate), method AACC 76-21.01, ICC 162.

### 2.10. Baking trials and texture analysis

The ingredients, based on 2000 g flour, 80 g fat, 40 g fresh yeast, 36 g salt, 30 g improver (E300, E471, E472e), 12 g calcium propionate and water as required. Dough was mixed in spiral mixer (Diosna, Osnabrück, Germany) for 3 min at a disk speed of 40 rpm and then for 7 min at a disk speed of 120 rpm. Water temperature was 20°C. Dough was rested in bulk for 5 min on the bench at room temperature, weighed into 1300-g portions, rounded manually, and rested further on the bench for 20 min at room temperature. Dough was then molded in a small-scale molder (Bertrand Puma, Portes-lès-Valence, France), placed into tins (L360 mm × D120 mm × H120 mm, open and closed) (Sasa, Le Cateau, France), and proofed at 38°C and 85% relative humidity for 50 min. Loaves were baked at 230°C top heat and 230°C bottom heat for 35 min in a deck oven (Picard, Canada). The oven was pre-injected with steam for 10 min. The loaves were depanned and allowed to cool for 120 min on cooling racks at room temperature. The loaves for analyses at day 7, 14 and 21 days were then packaged in containers and stored at 21°C.

For crumb texture analysis, bread was sliced transversely to obtain slices of 11-mm thickness. Two bread slices taken from the centre of each loaf were used to evaluate the crumb texture. Texture analysis was performed using a universal testing machine (TA-XT-Plus, Stable Micro Systems, Surrey, UK) equipped with a 25-kg load cell and a 35-mm aluminium cylindrical probe. The settings used were a test speed of 2 mm/sec with a trigger force of 20 g to compress the middle of the bread slice to 60% of its original thickness.

### 2.11. Starch characterization

Starch extraction and measurement of starch content were as described in Chia *et al*. (2017) with the following modifications. Starch was extracted from 100 grains and these were processed in a hand mill prior to grinding. CsCl and wash volumes were appropriately scaled. Starch content was measured using the Megazyme Total Starch Assay Kit (AA/AMG) (Megazyme International, Ireland) according to manufacturer’s instruction. Starch granule size distribution was determined using an automated image analysis system as above for developing grains (plot samples) or using a laser diffraction particle size analyser (Beckman Mastersizer 2000), according to the manufacturer’s instructions (pooled samples). Swelling power and amylose content were determined as described in Chia *et al*. (2107).

### 2.12. Statistical analysis

For Figure 4, data were modelled in R studio (version 4.0.4) using an analysis of variance, non-significant terms were dropped from the model, and then a TukeyHSD test was performed. For other Figures, statistical significance was tested using Student’s t-test in Excel. Probability (p) values below 0.05 were considered significant.

## 3. Results

### 3.1. Starch granule size and number in developing grain

Our analysis showed that the starch granule-size distributions of the wild types (two wild type sibling controls and the parent, Paragon) are very different from those of the three BlessT mutants (Fig. 1). B-granule number is very low, or absent, in BlessT grain throughout development. Up to 10 days after ear emergence, starch granule number per grain is similar in both wild types and mutants, and during this time the number of small, A-type granules decreases as their average size increases. In the mutants after 10 days, the number of granules per grain stabilises indicating that no more granules are initiated. The proportion of small granules continues to decrease as the A-granules grow. In the wild-types after 10 days, both the total number of granules and the number of small-granules increase as B-type granules are initiated. At 24 days, the grains reach maximum fresh weight and they begin to turn yellow (Supplementary Figure S1). Granule content per grain at 24 days is approximately equivalent to that at maturity (40 days), suggesting that granule initiation ceases as the grains enter the maturation phase. Before 24 days, the fresh weights of mutant and wild-type grain are very similar, however at maturity, the mutant grains have a slightly lower fresh weight.

**Figure 1.**
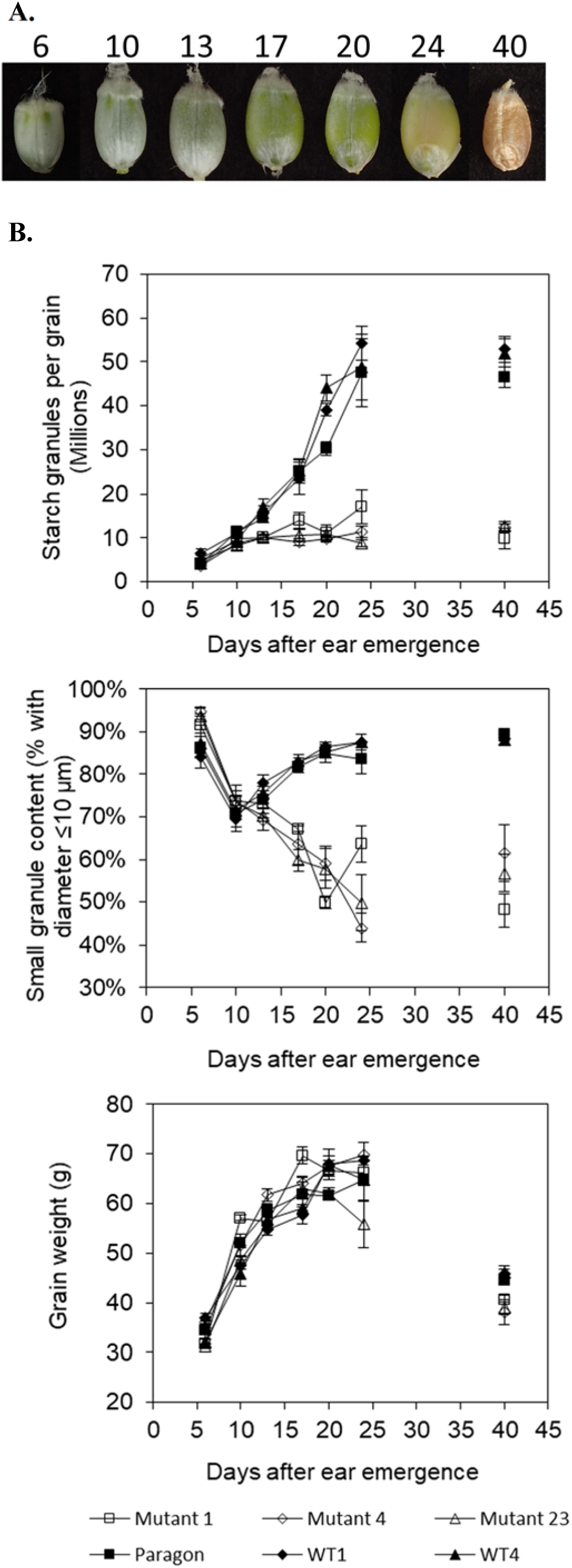
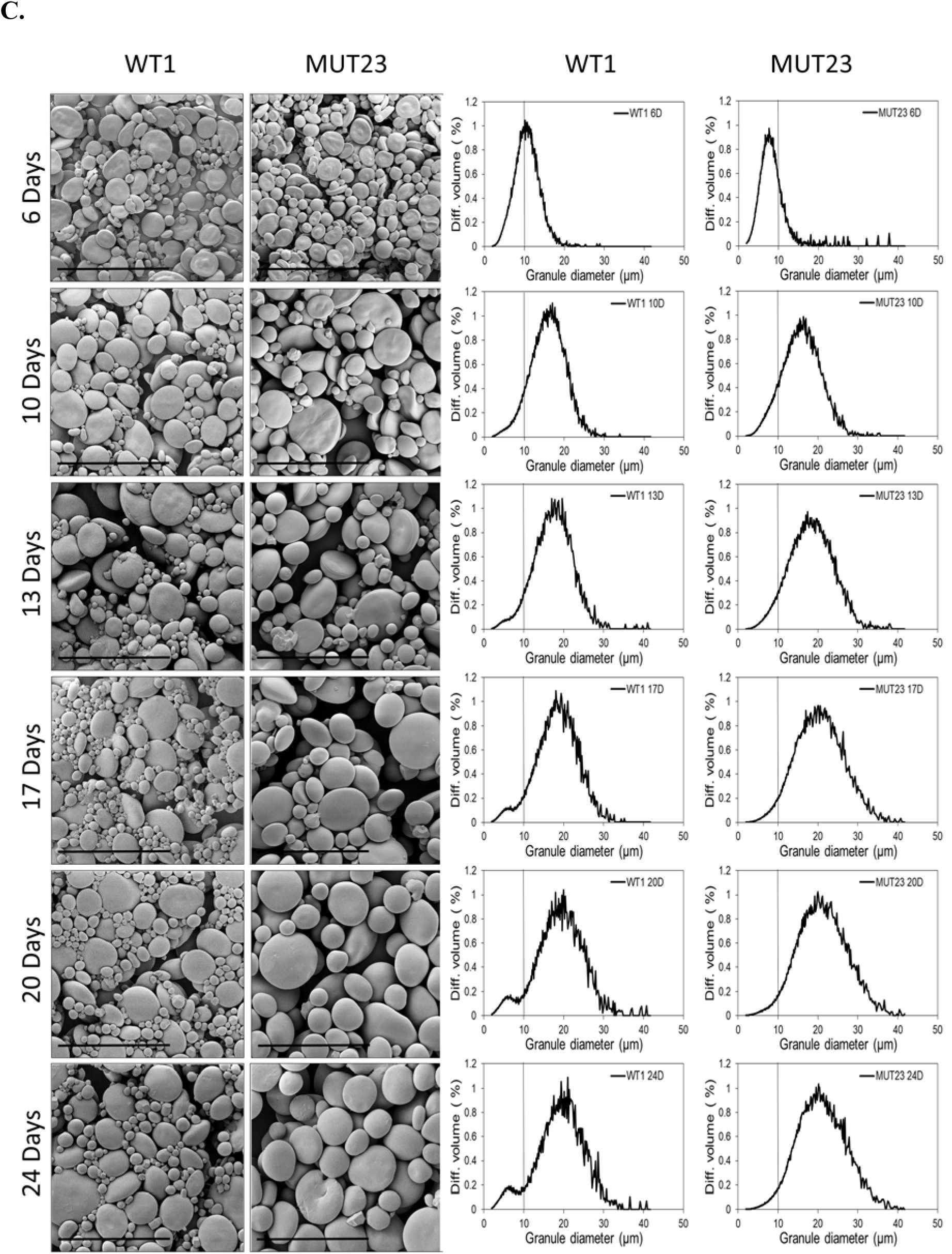
Starch granule size and number in developing grain. Grains were sampled from field-grown plants of a wild type (WT1) and a BlessT mutant (MUT23) at different stages of development. All grains were taken from the middle of the spike of a primary tiller. Images are for grains typical of each stage of development (values are days after ear emergence) (A). The fresh weights of individual developing and mature grains of Paragon (the latter were sampled at 40-41 days after ear emergence, Zadoks growth stage 59, and left to dry naturally before analysis) were determined (B). Starch extracts were prepared in triplicate. Samples comprised 6 or 12 grains from 3 or 6 different plants. An automated image analysis system was used to determine the total number of starch granules per endosperm and the number of ‘small’ granules (with diameters ≤10 μm) per endosperm. Depending on the stage of development and genotype, the ‘small’ granule category can include B-type granules and/or small A-type granules. On all graphs, values are means ± SE. Markers are unfilled for the three mutants and filled for the wild-types. MUT1 and Paragon markers are squares, MUT4 and WT1 markers are diamonds and MUT23 and WT4 markers are triangles. Starch granule-size distribution was assessed qualitatively using SEM imaging and quantitatively using a Coulter Counter (C). Data and images for developing grains are shown. The scale bar on the SEM micrographs is 50 μm. Coulter counter values are means for 3 replicate starch preps and for guidance a line is drawn on each graph at 10 μm diameter..

### 3.2. Growth metrics

For plants grown in field plots in 2018, the following five plant growth metrics were measured: plant establishment, plant density, tiller number, flowering time and plant height (Fig. 2, Supplementary Table S1). No consistent differences between all the three mutants and both wild type sibling controls were found for any of these parameters (Student’s t-test; p value >0.05). This suggests that the lack of *BGC1-4A* and *BGC1-4D* does not cause plant growth defects.

**Figure 2.**
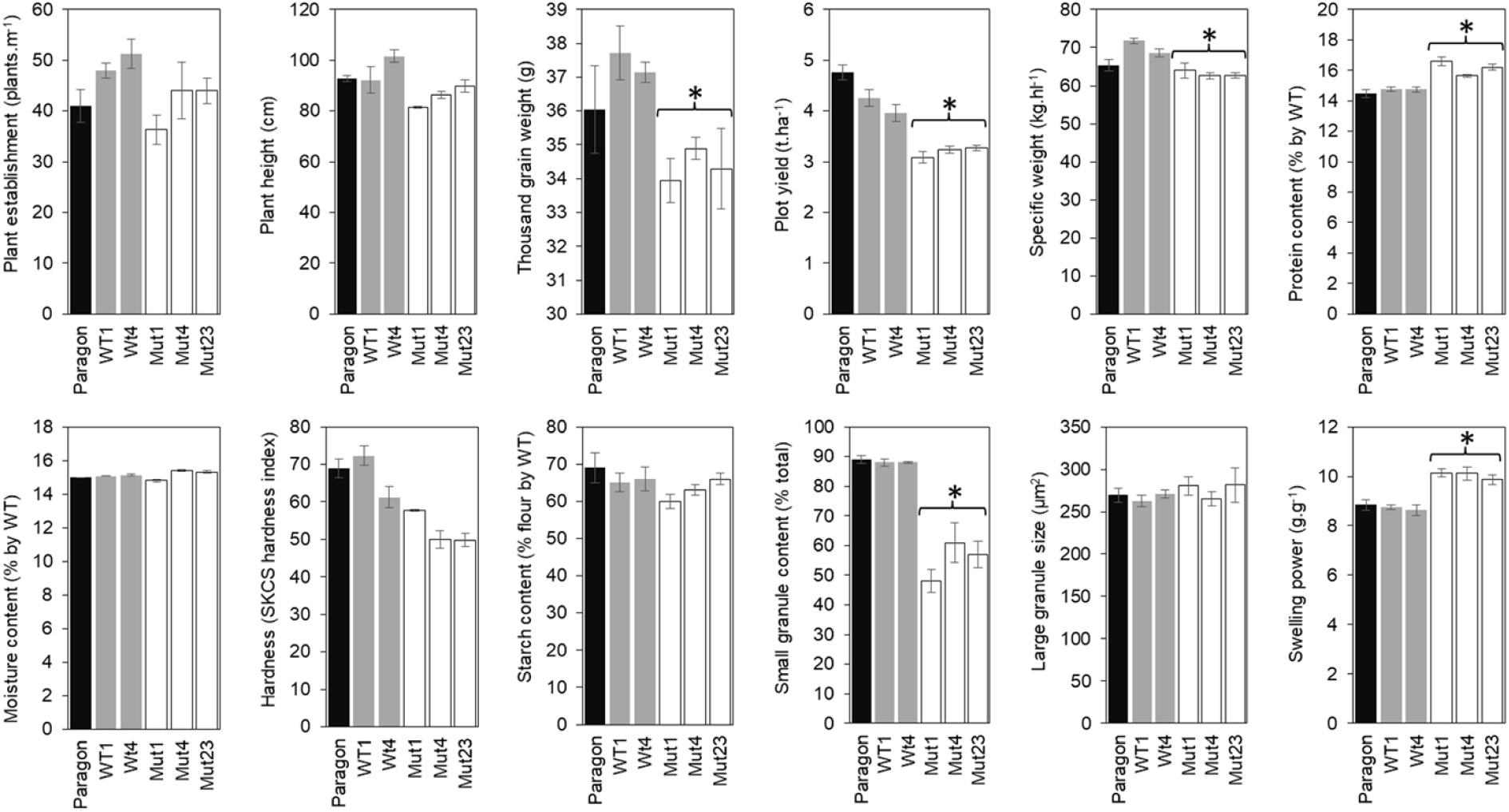
Characterization of plants, grain, flour and starch. Plants, grains and flours from B-granule-less deletion mutant lines, wild-type sibling control lines and Paragon, the parental cultivar were characterized. Values are means ± standard errors. MUT = mutant, WT = wild type. Stars indicate values that are significantly different in pair-wise comparisons of all three MUTs with both WT controls. Selected characterization data are shown. For the full data set, replication details and all pair-wise statistical analysis see Supplementary Table 1. Data are for field-grown wheat from 2018.

Some differences were observed for individual MUT or WT lines: plant establishment was reduced in MUT1 compared with the WTs, plant height was increased in WT4 compared to the MUTs, WT1 flowered later than Paragon and WT4 was taller than Paragon. These effects on individual lines may be due to the deletion of genes unlinked to *BGC1* that vary between these lines.

### 3.3. Yield parameters

The following six yield parameters were measured: thousand grain weight, grain area, grain width, grain length, plot yield and hectolitre weight (Fig. 2, Supplementary Table S1). Of these, three parameters (thousand grain weight, plot yield and specific weight) were decreased in the MUTs compared to the WTs. Analysis of the pooled samples also showed that BlessT grains had a lower specific weight and thousand grain weight than WT controls (Supplementary Table S2). However, plot yield was decreased in the WTs and MUTs relative to Paragon, and specific weight was decreased in WT1 relative to Paragon. This suggests that genes other than *BGC1* may be responsible for, or contribute to, the lower yields in the deletion lines.

### 3.4. Grain and flour characteristics

The following seven characteristics were measured: starch content, protein content, moisture content, hardness, α-amylase activity, germination and viability (Fig. 2, Supplementary Table S1). Of these, only the protein content was consistently different between the MUTS and the WTs, whilst moisture content and grain hardness were statistically significantly different for some of the MUT/WT comparisons only. The protein content in the MUTs was ∼10% higher than that in the WTs, whilst the protein content of the WTs and Paragon were not statistically significantly different. The moisture contents of WT1 were lower than those of all of three MUTs whilst that of WT4 was lower than MUT1 and MUT4. The moisture contents of both WTs was greater than that of Paragon. With respect to grain hardness, WT1 grains were harder than all MUT grains whilst WT4 grains were harder than those of MUT4 and MUT23. There was no difference in hardness between the WTs and Paragon. Analysis of the pooled grain samples also showed that BlessT grains had a higher protein content and that their endosperm was softer and that ash content was also higher (Supplementary Table S2). Together these results suggest that increased protein, ash and moisture content, and decreased grain hardness are associated with deletion of *BGC1*.

### 3.5. Amylase activity

Alpha amylase activity was measured in three ways: by direct enzymatic assay, as Hagberg falling number and from RVA analysis (Fig. 3, Supplementary Table S1, Supplementary Table S2). For the 2018 individual plot and pooled grain samples, there were no differences between the values for the MUTs and WTs, or between the WTs and Paragon. However, these amylase activities were generally high (low Hagberg falling numbers) and high preharvest amylase production was observed generally for wheat in the Cambridge area in 2018, in which there was a higher-than-average rainfall. For this reason, we also measured the α- amylase activity of samples from plants grown in a glasshouse in 2017. In this environment, the α-amylase activity in Paragon and the WTs was reduced relative to the field grown grains but it was increased in the MUTs. Thus, the lack of B-type granules appears, in some environments, to be associated with increased activity of α-amylase.

**Figure 3.**
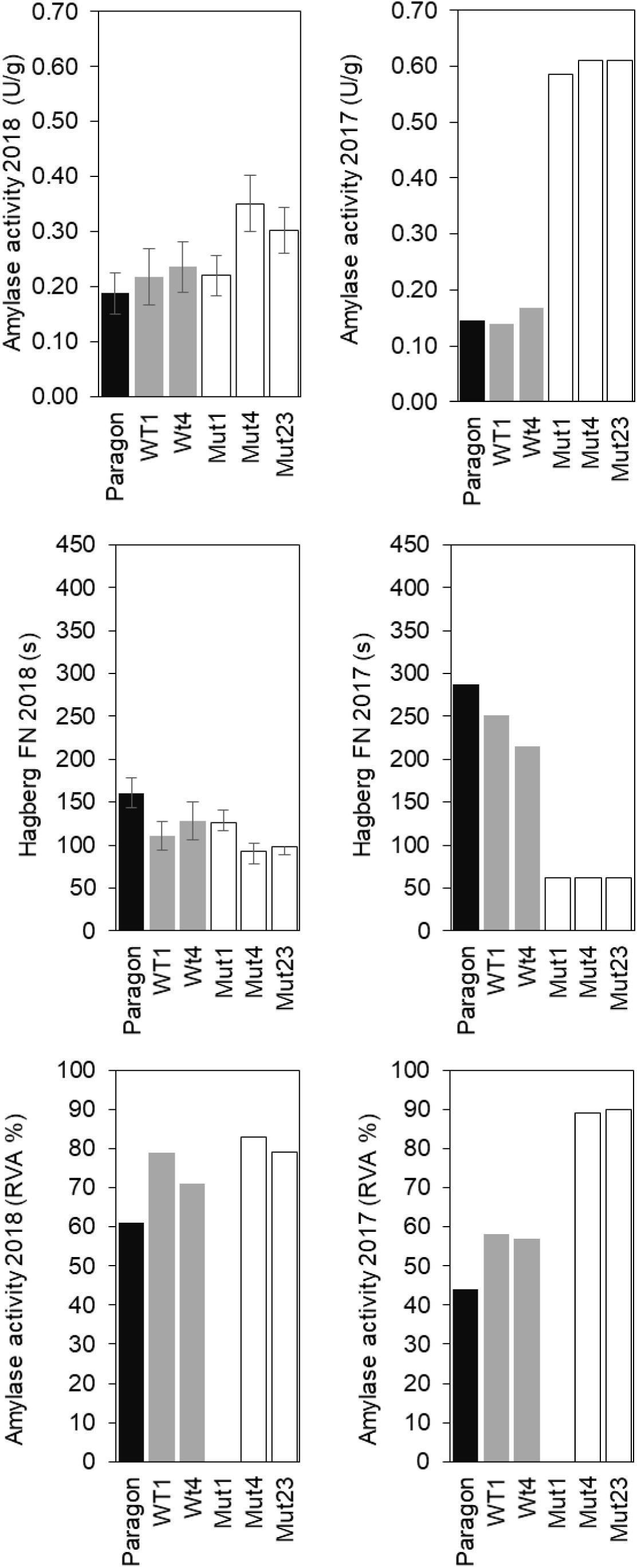
Comparison of α-amylase activity in mature grains. The activity of α-amylase in milled flour from mature grains was determined using three different methods: direct enzymatic assay, Hagberg falling number (HFN) and indirect RVA estimation. For the enzymatic determination and HFN of samples from 2018, values are means ± SE for four biological replicates. For all other measurements, values are means of duplicate technical replicates only and no errors are shown. Wheat was grown in the soil in a glasshouse in 2017 or in the field in 2018.

### 3.6. Starch characterization

The following five starch function properties were measured: small granule content, small granule size, large granule size, swelling power and amylose content (Fig 2, Supplementary Table S1). Of these, only the small granule content and the swelling power was consistently different between the MUTS and the WTs, the small-granule content being lower and the swelling power higher in the MUTs. The reduction in small-granule content in mature grain is consistent with the lack of B-type granules in BlessT observed in developing grains (Fig. 1).

### 3.7. Milling and baking characteristics

Due to the quantities of grains required for these tests, they were carried out on the pooled-grain samples only. There was no difference between BlessT and wildtype in milling yield. However, the level of starch damage in the MUTs was lower than that in the WTs or Paragon, which is consistent with reduced grain hardness. The rheological characteristics of the flours were compared by Glutomatic, Alveograph and RVA. These suggested that MUT flour has good gluten quality, with the wet and dry gluten values correlating with the increased protein content. The separability of starch and gluten was also good in the MUTs. The dough Elasticity/Extensibility (P/L) ratio was higher in the MUTs, probably due to a lack of hydration: the Farinograph analysis showed that MUT flour required a higher level of hydration than WT or Paragon flour. The Farinograph also showed that MUT dough had a poor overall stability (which may be related to the moderately high α-amylase activity in all genotypes). Using standard conditions, all RVA values were low. This is consistent with the high α-amylase activity observed in these flours and was tested by including silver nitrate in the samples to inhibit amylase activity. In the presence of silver nitrate, the RVA values for all flours were increased and no differences were detected between genotypes.

Baking trials were conducted on MUT, WT, Paragon and an internal LCI control flour (Fig. 4, Supplementary Table S3). Acceptable loaves were produced indicating that the high α-amylase activity in the test flours had little or no detrimental impact (Fig. 4A). Dough making confirmed that the MUT flour required more water than the controls (58% vs 54% w/w, respectively). Even so, the MUT loaves were under-developed (suggesting that even more water would have been optimal).

**Figure 4.**
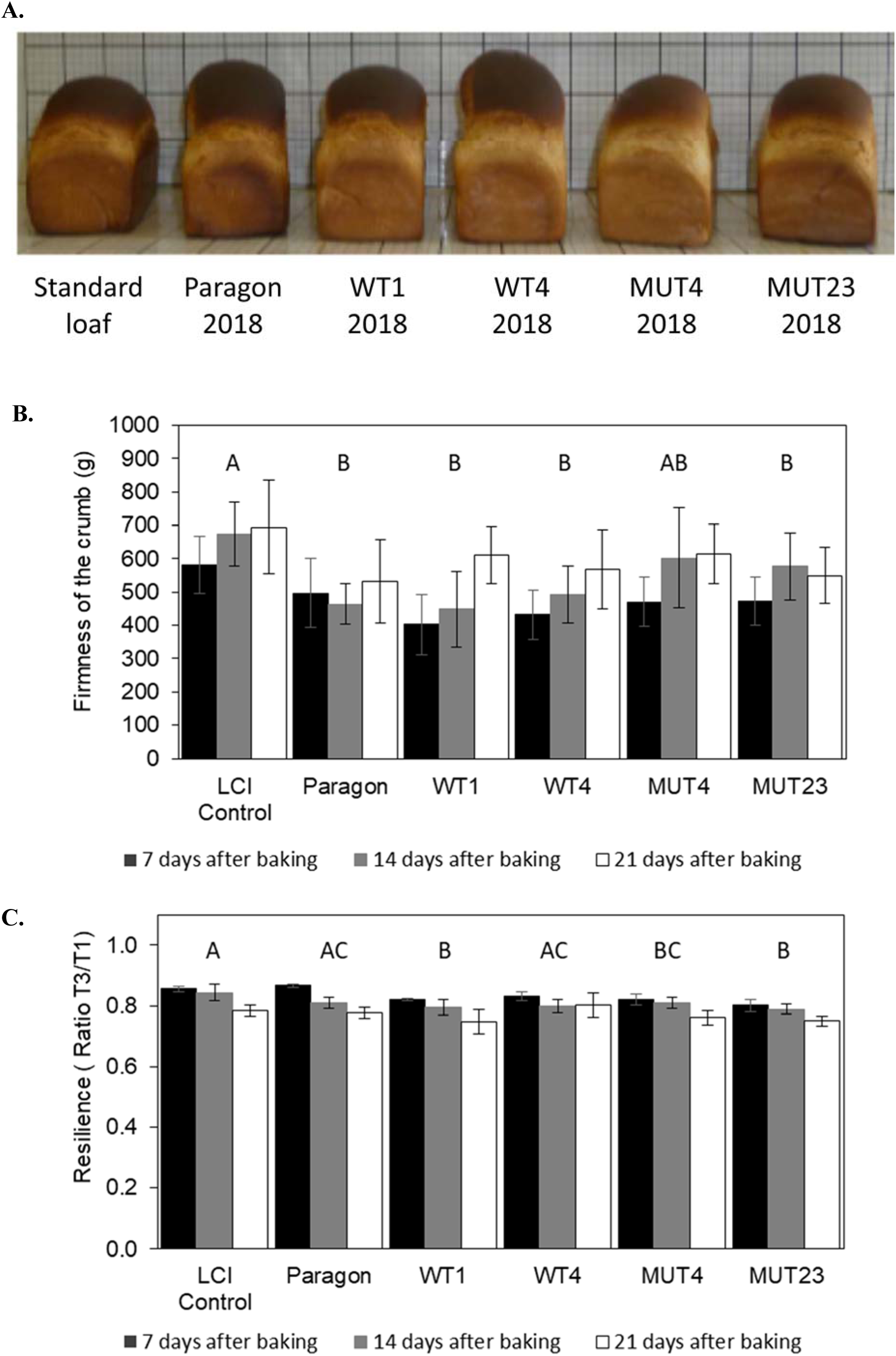
Baking trials. Flour was prepared from the pooled grains grown in the field in 2018. The water content of the dough was adjusted to give a suitable consistancy. Loaves were baked using both closed (not shown) and open molds (A). The crumb structure of slices of loaves was compared after 7,14 and 21 days storage using a texture analyser. For determination of firmness (B), the force of resistance developed by the slice bread against penetration by the probe is determined. In the determination of resilience (C), a double compression was used. The resilience value is the ratio of the area under curve for the second compression relative to that for the first compression. Statistical analyses (B,C) were done using TukeyHSD and significant differences (p<0.05) between genotypes (over all timepoints) are indicated by different letters.

In the texture trials, all genotypes showed a tendency to increased firmness with time in storage (Fig. 4B). Statistical analysis showed significant differences between 7-14 days, (p=0.03) and 7-21 days (p ≤0.001) but not between 14-21 days (p=0.12). An increase in firmness with time is expected since staling makes the crumb structure of the internal matrix stiffer (firmer) and more brittle (crumbly). For these data, the interaction between age and genotype was not significant (p=0.505) and there were no significant differences between genotypes at any given time point (p=0.505). However, there were significant differences (p< 0.05) between genotypes over all timepoints, as indicated by different letters in Figure 4B. Bread made from the LCI control flour was firmer than all of the other breads but there was no difference in firmness between the Paragon, WT and MUT breads.

Resilience is correlated with the extent of staling. Typically, the fresher the bread, the higher is its resilience, whilst stale bread shows little or no resilience. As expected, for all genotypes the resilience of bread reduced with time in storage with each comparison being significant (p ≤0.001) (Fig. 4C). There was a significant interaction between age and genotype for these data and the complete statistical analysis of differences within age groups for each genotype is shown in Supplementary Table S3(B). There were significant differences (p< 0.05) between genotypes (over all timepoints) as indicated by different letters in Figure 4C. However, there were no consistent differences between the WTs and MUTs suggesting that the lack of B-granules did not affect resilience.

Overall these studies suggest that the lack of B-type starch granules has no major detrimental impact on bread making or bread texture: with respect to baking properties, dough behaviour and crumb structure, the MUTs were comparable to the WTs. However, these are preliminary studies due to the limited amounts of grain available and further in-depth analysis will be needed.

## 4. Discussion

BlessT wheat has several unique functional properties including higher water absorption of starch and flour, higher protein content (BlessT has 110% by dry weight protein content compared to that of the wild type sibling controls) and reduced grain hardness. Unlike most mutations affecting starch synthesis which result in lower starch content, shrivelled grain and substantially reduced yield, BlessT has a modest, if any, reduction in grain weight. This, together with normal starch content in flour, suggests that it should be possible (using for example, TILLING mutants or CRISPR) to produce commercial wheat varieties that lack B-type starch granules without compromising yield. These findings support the potential utility of this novel type of wheat as a specialist crop in applications ranging from bread making and alcohol production to improved industrial starch products.

There is some evidence of reduced grain hardness in BlessT flour. The main factor controlling grain hardness in wheat is puroindoline content (Morris, 2002). We do not yet know whether the puroindoline content is altered in BlessT wheat but it is possible that the change in granule-size distribution itself is directly responsible for the effect on hardness observed. Although modest, the reduction in hardness in BlessT is accompanied by a reduced level of starch damage after milling. Despite this and its other unique properties, BlessT flour was found to make bread with acceptable characteristics although more water needed to be added to form bread dough of suitable consistency.

The main detrimental effects of BlessT compared to wild type sibling controls were reduced yield (plot yield and specific weight) and variable, sometimes high, levels of α- amylase. The yield parameters of the wild type sibling controls were also reduced relative to Paragon, suggesting that deletions of genes other than *BGC1* in both the wild type siblings and the BlessT mutants may be responsible for reduced yield. In support of this, starch content per grain was not different between BlessT and the wild type sibling controls. The effects of reduced *BGC1* in BlessT on α-amylase levels were complex. The levels of α-amylase in BlessT were higher than normal when grains were grown in a glasshouse but no different from normal when grains were grown in a field in a particularly wet year. At present we do not know to what extent these effects on α-amylase are caused by genes other than *BGC1*. This is currently being tested by creating TILLING mutant lines of hexaploid wheat with reduction in *BGC1* specifically.

The loss of B-granules in BlessT is partially compensated for by an increase in starch in A-granules. B-type starch granules account for a smaller proportion of the total granule volume than A-type granules in mature wild type grains e.g. 16% of starch volume is B-granules in mature grains of WT1 (Coulter counter data, not shown). However, BlessT mutants do not have substantially reduced grain size (the thousand grain weight of the mutants was reduced by less than 10% relative to the wild type sibling controls and was not different from that of Paragon) and neither starch content (as % flour) nor milling yield were affected (Supplementary Table S1, Supplementary Table S2). This suggests that there is some diversion of starch deposition into the A-granule fraction in BlessT.

There have been many attempts to compare the characteristics of wheat A- and B-type starch granules and these have often shown variations between granule types. For example, both the amylose content and the gelatinization enthalpy of B-type starch is higher than that of A-granules (Chiotelli and Le Meste, 2002; Wei *et al*., 2010, Li *et al*., 2013). However, the observed variations between granule types have not always been consistent between studies. For example, the swelling power of B-granules was found to be greater than that of A-type granules (Chiotelli and Le Meste, 2002; Wei *et al*., 2010) or less than that of A-type granules (Li *et al*., 2013). Since BlessT starch has higher swelling power than wild type starch in our study, this could indicate that the swelling power of B-granules is less than that of A-type granules. However, given the possibility that the A-type granules in BlessT are different from those in wild type wheat, this conclusion is presently unsafe: the A-granules in BlessT could swell more than the A-granules in wild type wheat.

BlessT wheat was made using deletion mutants and it is affected in many genes, not just the gene responsible for controlling B-granule content, *BGC1*. In this study, we have attempted, by using multiple mutant and control lines, to explore the likely effects of B-granule elimination on the characteristics of wheat. However, we are also attempting to produce BlessT#2 hexaploid wheat by combining TILLING mutant lines lacking specific *BGC1* genes but with few other genetic alterations. We have already established using TILLING mutants of tetraploid wheat that reduction in the level of *BGC1* to within a critical range is required to achieve elimination of B-type starch granules without impacting on the production of A-type granules (Chia *et al*., 2020). Our hypothesis is that hexaploid TILLING mutants lacking the *BGC1-4A* and *-4D* homeologues of *BGC1*, but having a normal *BGC1-4B* homoeologue will lack B-type granules. Such B-granule-less TILLING lines will allow us to test which of the characteristics of BlessT observed here are attributable to *BGC1* rather than to other, background gene modifications, and will provide lines for breeding and further functionality testing.

## Abbreviations

BGC1: B-granule content
DWT: Dry weight
HFN: Hagberg falling number
NIR: near-infrared
NDVI: Normalized Difference Vegetation Index
RVA: rapid visco analysis.

## Supplementary data

**Supplementary Table S1.**
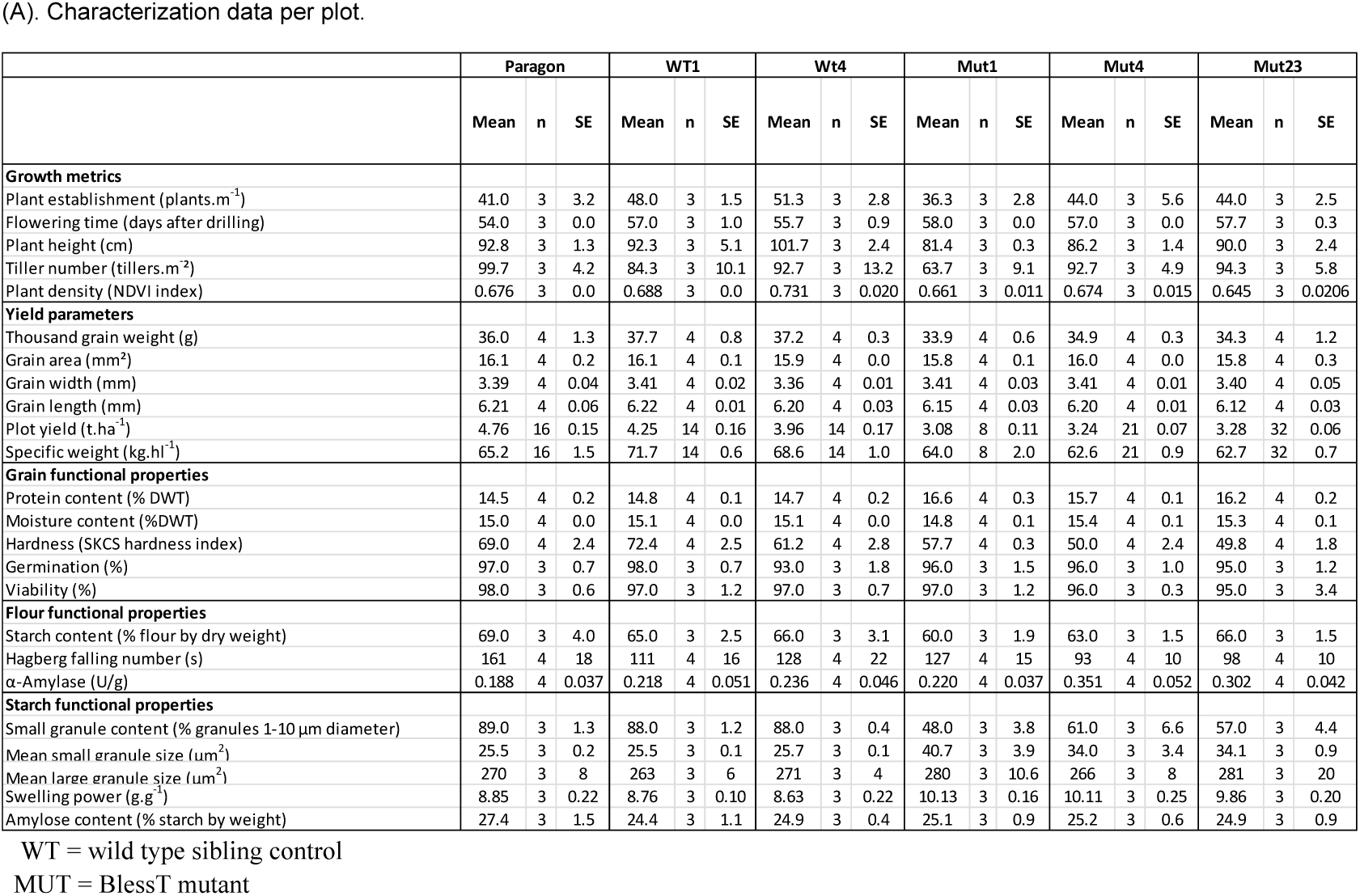
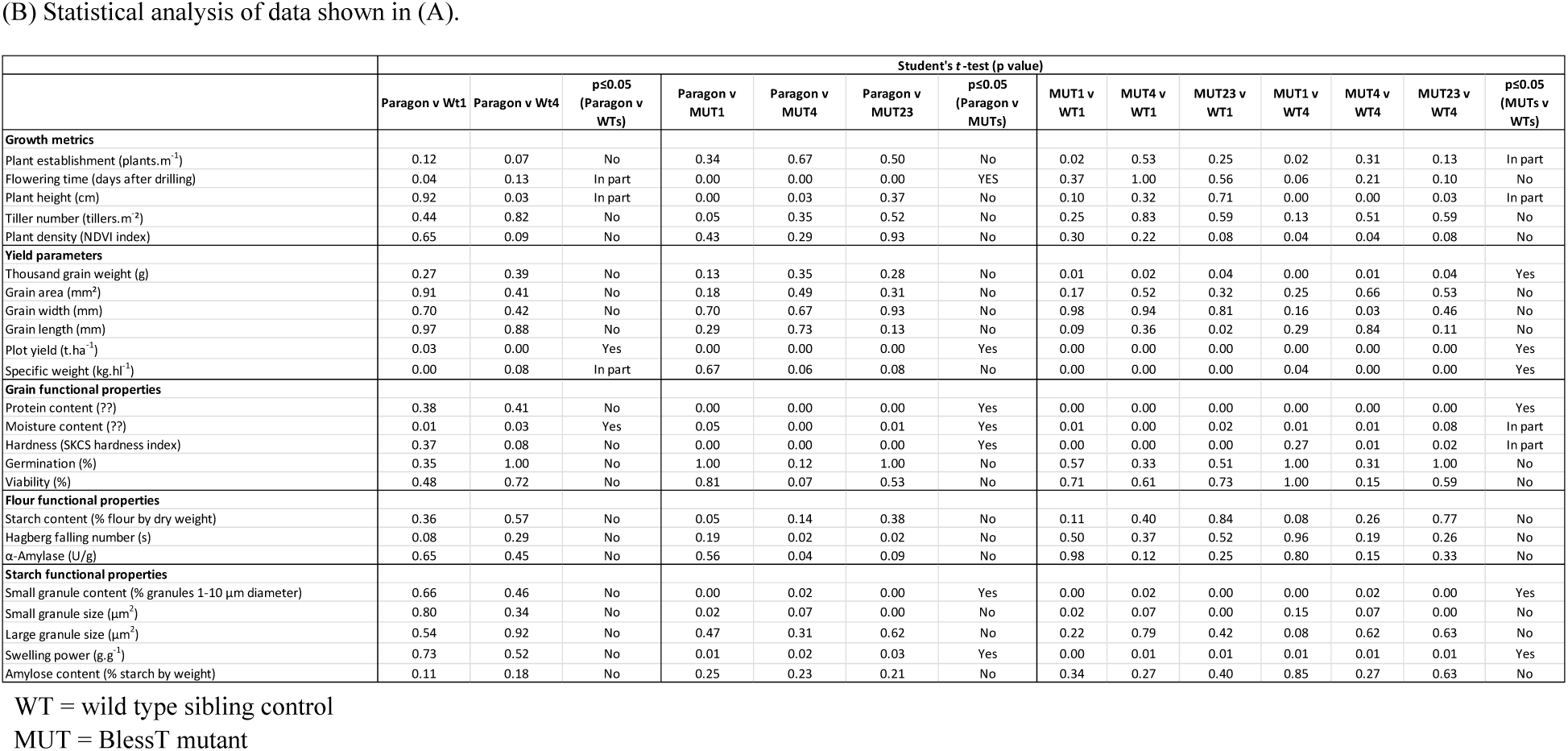
Characterization data for plants grown in field plots in 2018.

**Supplementary Table S2.**
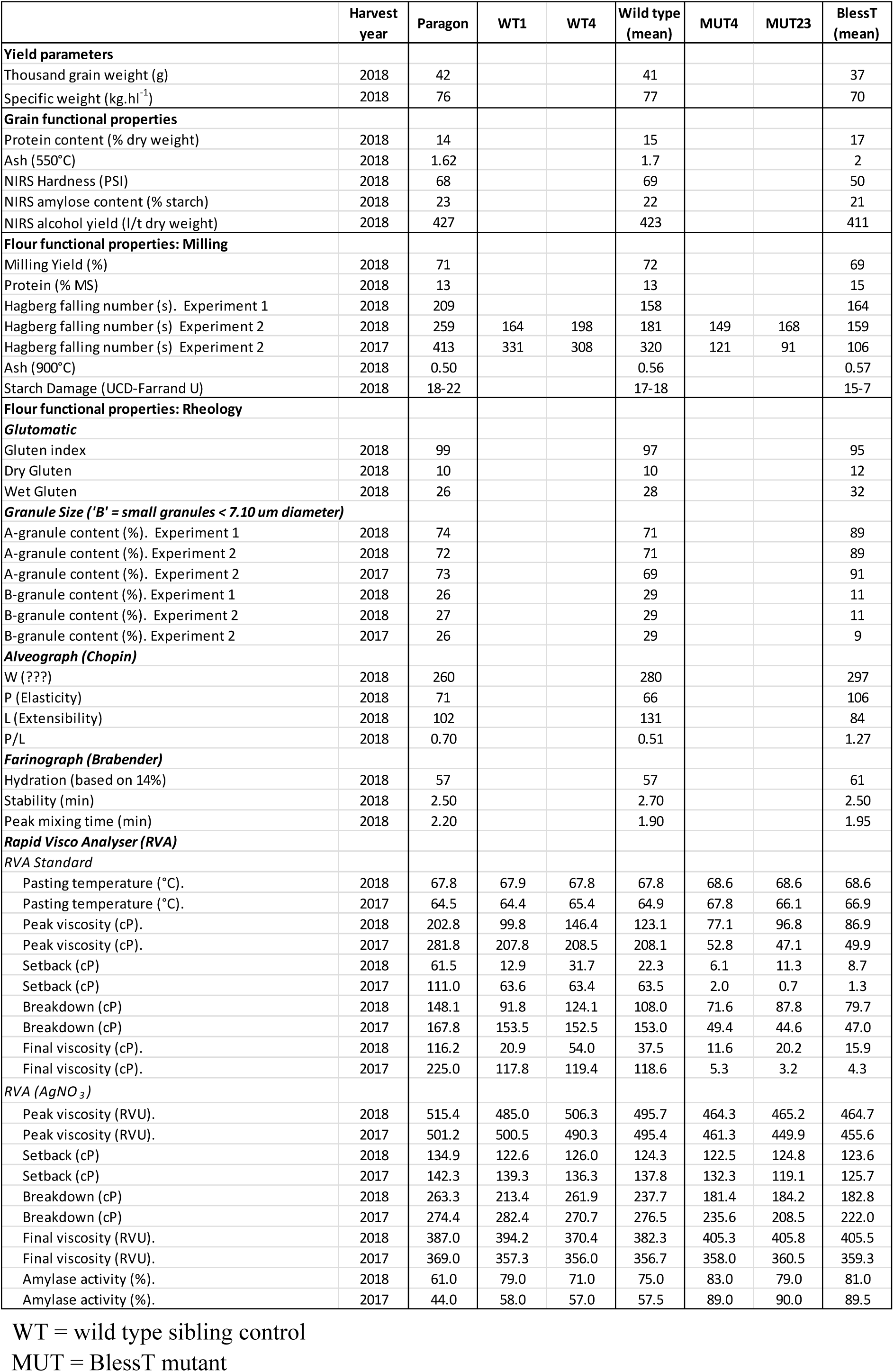
Characterization data for pooled grain samples from 2018.

**Supplementary Table S3.** Characterization of bread baking quality.

(A). Values for firmness and evolution of resilience
|  | Days after baking | LCI Control | Paragon | WT1 | WT4 | WT (mean) | MUT4 | MUT23 | MUT (mean) |
| --- | --- | --- | --- | --- | --- | --- | --- | --- | --- |
| **Baking quality** |  |  |  |  |  |  |  |  |  |
| Firmness of the crumb (g) | 7 | 581 ± 87 | 496 ± 104 | 402 ± 91 | 432 ± 75 | 418 | 471 ± 73 | 472 ± 74 | 472 |
| (Resistant force developed by the sliced bread against probe penetration). | 14 | 673 ± 96 | 463 ± 62 | 448 ± 114 | 492 ± 84 | 470 | 602 ± 151 | 577 ± 100 | 590 |
|  | 21 | 694 ± 139 | 532 ± 125 | 611 ± 85 | 567 ± 119 | 590 | 614 ± 90 | 549 ± 84 | 582 |
| Evolution of resilience | 7 | 0.86 ± 0.01 | 0.87 ± 0.01 | 0.82 ± 0.01 | 0.82 ± 0.01 | 0.83 | 0.82 ± 0.02 | 0.80 ± 0.02 | 0.81 |
| (Ratio of resistant forces, peak 2nd test : peak 1st test. A value of 1 indicates maximum resilience). | 14 | 0.84 ± 0.03 | 0.81 ± 0.02 | 0.80 ± 0.03 | 0.80 ± 0.02 | 0.80 | 0.81 ± 0.02 | 0.79 ± 0.02 | 0.80 |
|  | 21 | 0.78 ± 0.02 | 0.78 ± 0.02 | 0.04 | 0.75 ± 0.04 | 0.78 | 0.76 ± 0.02 | 0.75 ± 0.02 | 0.76 |

| Genotype | Days after baking |  |  |
| --- | --- | --- | --- |
|  | 7 | 14 | 21 |
| LCI control | A | A | AB |
| Paragon | A | AB | AB |
| WT1 | AB | AB | A |
| WT4 | AB | AB | B |
| MUT4 | AB | AB | AB |
| MUT23 | B | B | A |

**Supplementary Figure S1.**
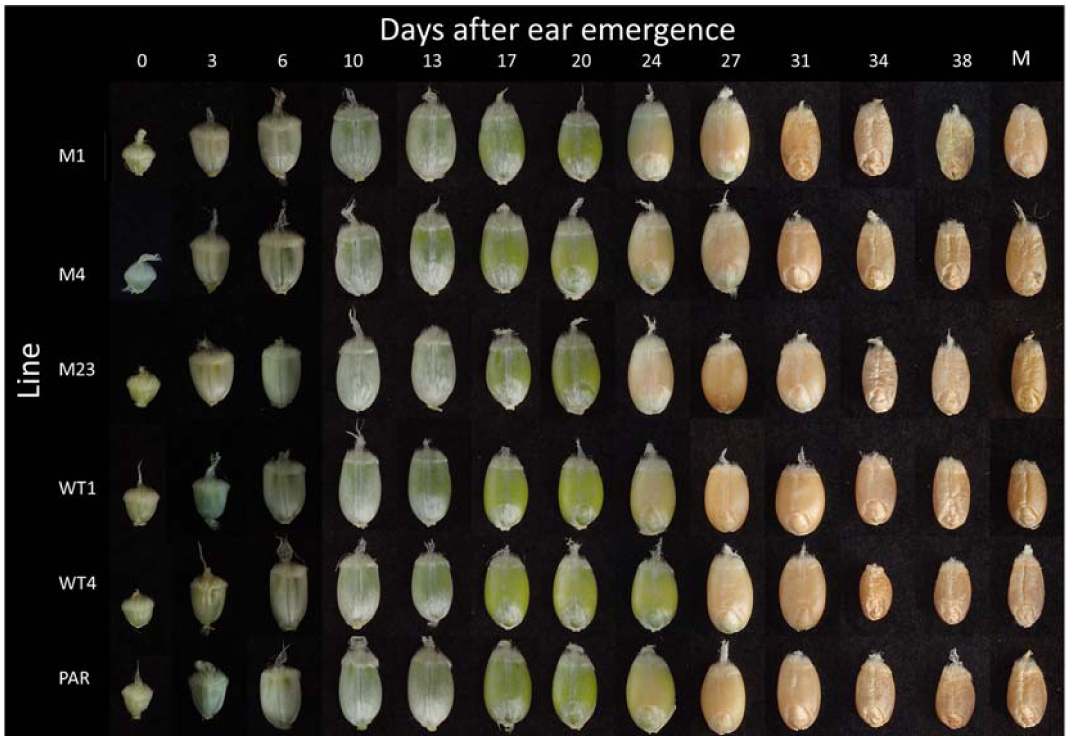
A montage of images of developing grains. M1, M4 and M23 are BlessT mutant lines, WT1 and WT4 are wildtype sibling lines and PAR is the parental cultivar, Paragon. M= mature grains harvested at 40 or 41 days after ear emergence. Typical images are shown.

**Supplementary Figure S2.**
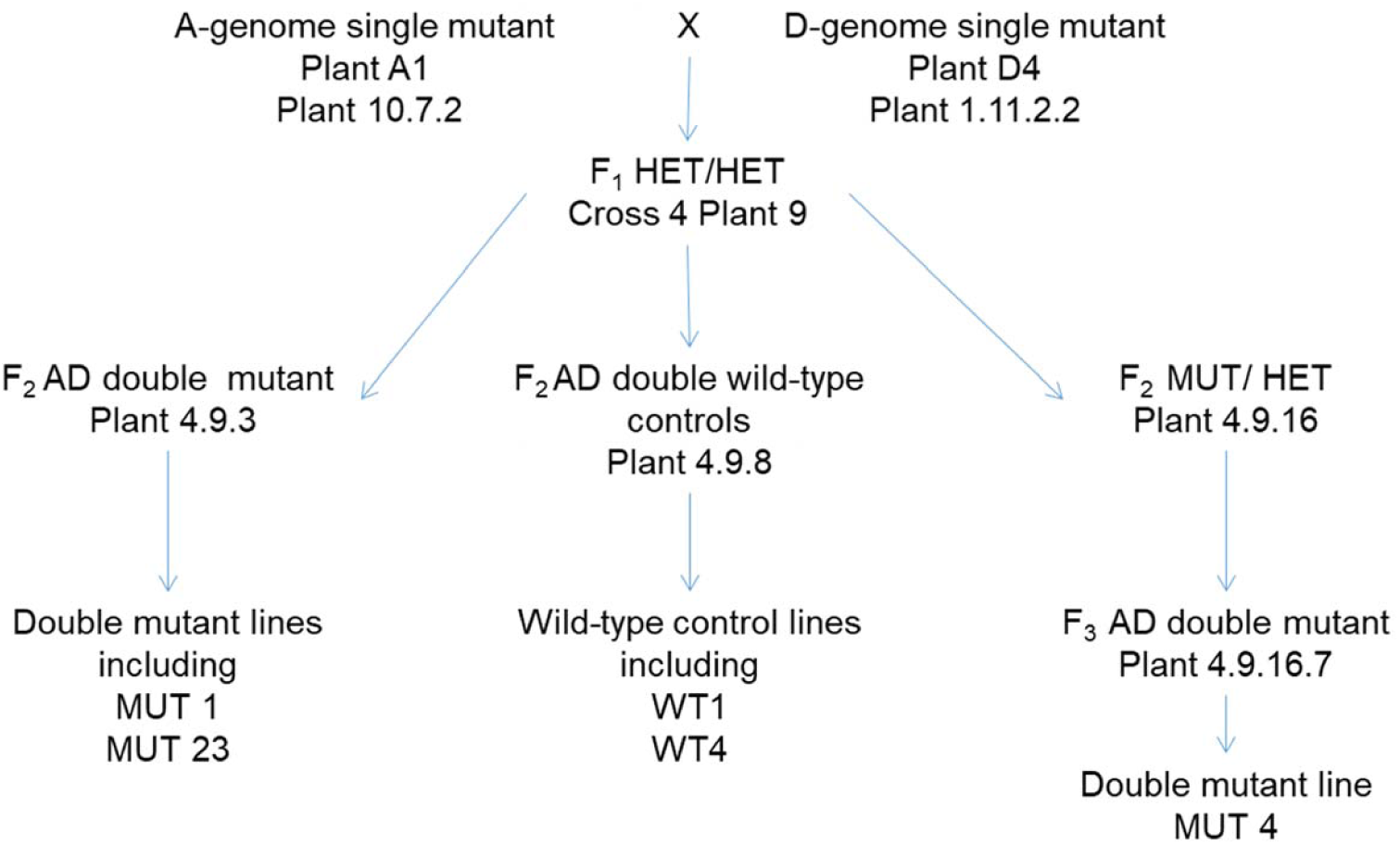
The pedigree of the BlessT lines and controls.

